# ALLSorts: a RNA-Seq classifier for B-Cell Acute Lymphoblastic Leukemia

**DOI:** 10.1101/2021.08.01.454393

**Authors:** Breon Schmidt, Lauren M. Brown, Georgina L. Ryland, Andrew Lonsdale, Hansen J. Kosasih, Louise E. Ludlow, Ian J. Majewski, Piers Blombery, Paul G. Ekert, Nadia M. Davidson, Alicia Oshlack

**Affiliations:** Peter MacCallum Cancer Centre, Parkville, VIC, Australia; School of BioScience, University of Melbourne, VIC, Australia; Murdoch Children’s Research Institute, Parkville, VIC, Australia; Children’s Cancer Institute, Lowy Cancer Centre, UNSW Sydney, Kensington, NSW, Australia; Sir Peter MacCallum Department of Oncology, University of Melbourne; The Walter and Eliza Hall Institute of Medical Research, 1G Royal Parade VIC 3052 Australia; University of Melbourne, Department of Medical Biology, 1G Royal Parade VIC 3052 Australia; Centre for Cancer Research, University of Melbourne, VIC, Australia; Department of Paediatrics, University of Melbourne, Parkville, VIC, Australia; School of Women’s and Children’s Health, UNSW Sydney, Sydney, NSW, Australia; University of New South Wales Centre for Childhood Cancer Research, UNSW Sydney, Sydney, NSW, Australia

**Author notes:** Corresponding Author: Alicia Oshlack.

## Abstract

B-cell acute lymphoblastic leukemia (B-ALL) is the most common childhood cancer. Subtypes within B-ALL are distinguished by characteristic structural variants and mutations, which in some instances strongly correlate with responses to treatment. The World Health Organisation (WHO) recognises seven distinct classifications, or *subtypes*, as of 2016. However, recent studies have demonstrated that B-ALL can be segmented into 23 subtypes based on a combination of genomic features and gene expression profiles. A method to identify a patient’s subtype would have clear clinical utility. Despite this, no publically available classification methods using RNA-Seq exist for this purpose.

Here we present ALLSorts: a publicly available method that uses RNA-Seq data to classify B-ALL samples to 18 known subtypes and five meta-subtypes. ALLSorts is the result of a hierarchical supervised machine learning algorithm applied to a training set of 1223 B-ALL samples aggregated from multiple cohorts. Validation revealed that ALLSorts can accurately attribute samples to subtypes and can attribute multiple subtypes to a sample. Furthermore, when applied to both paediatric and adult cohorts, ALLSorts was able to classify previously undefined samples into subtypes.

ALLSorts is available and documented on GitHub (https://github.com/Oshlack/AllSorts/).

**Key Points:**

- ALLSorts is a gene expression classifier for B-cell acute lymphoblastic leukemia, which predicts 18 distinct genomic subtypes - including those designated by the World Health Organisation (WHO) and provisional entities.
- Trained and validated on over 2300 B-ALL samples, representing each subtype and a variety of clinical features.
- Correctly identified subtypes in 91% of cases in a held-out dataset and between 82-93% across a newly combined cohort of paediatric and adult samples.
- ALLSorts assigned subtypes to samples with previously unknown driver events.

ALLsorts is an accurate, comprehensive and freely available classification tool that distinguishes subtypes of B-cell acute lymphoblastic leukemia from RNA-sequencing.

## Introduction

B-cell acute lymphocytic leukemia (B-ALL) is the most common childhood malignancy and is a rare Leukemia in adults (Gu et al., 2019; Hunger & Mullighan, 2015; Inaba, Greaves, & Mullighan, 2013; Terwilliger & Abdul-Hay, 2017). B-ALL subtypes are distinguished by characteristic structural variants and mutations, which in some instances strongly correlate with responses to treatment (Gu et al., 2019; Inaba et al., 2013; Paietta et al., 2021; Terwilliger & Abdul-Hay, 2017). The 2016 World Health Organisation (WHO) classification recognises seven distinct B-ALL subtypes (Arber et al., 2016). These are defined by either the presence of a fusion gene (*BCR-ABL1, TCF3-PBX1, ETV6-RUNX1, IGH-IL3*, and *KMT2A* rearrangements in KMT2A) or chromosomal aneuploidy (hyperdiploidy and hypodiploidy). There are two provisional subtypes, intrachromosomal amplification of chromosome 21 (iAMP21) and Philadelphia-like (Ph-like) (Arber et al., 2016). iAMP21 is relatively rare (~2% of cases) and represents a complex structural change within chromosome 21, including amplification of a region including the *RUNX1*, *ETS* and *ERG* genes (Inaba et al., 2013; Tsuchiya, Davis, & Gardner, 2017). Ph-like has a gene expression profile that resembles Ph positive (*BCR-ABL1*) B-ALL but is driven by activating mutations in other kinases.

The WHO classification of B-ALL is incomplete. Recognised driver mutations are not identified in all cases. In addition, combinations of unsupervised and supervised machine learning suggest the existence of up to 23 subtypes (Gu et al., 2019; Lilljebjörn et al., 2016). Although it remains to be validated, subtype assignment has the potential of extending and refining the current standards of risk stratification. Indeed, the current standard of care already incorporates some molecular classification in order to identify patients at higher risk of disease relapse (Inaba, Azzato, & Mullighan, 2017; Schultz et al., 2007). Detection of *BCR-ABL1* indicates high-risk disease and treatment should be modified to include a ABL1-targeting tyrosine kinase inhibitor such as imatinib (Inaba et al., 2013). *ETV6-RUNX1* fusions indicate a much lower risk of treatment relapse (dependent on some clinical variables) (Brown et al., 2020; Inaba et al., 2017; Schultz et al., 2007). More recently, next generation sequencing of RNA (RNA-Seq) has been used to identify fusion genes, quantify gene expression, and perform variant calling to identify a larger number of drivers (Brown et al., 2020; Byron, Van Keuren-Jensen, Engelthaler, Carpten, & Craig, 2016), and is making its way into diagnostic pipelines (Brown et al., 2020; Inaba et al., 2017). Although gene expression quantification is particularly useful for identifying molecular subtypes, there is currently no publicly available software for taking RNA-Seq expression and performing this classification.

Here we present ALLSorts: a B-ALL gene expression classifier that attributes samples to 18 subtypes previously defined by Gu et al. (2019). ALLSorts has several unique features including a hierarchical design that offers broader group classifications if more specific subtypes cannot be ascertained. ALLSorts also incorporates custom features guided by B-ALL biology as suggested by Lilljebjörn et al. (2016). Additionally, ALLSorts can attribute multiple subtypes to samples - the utility of which was demonstrated by Nordlund et al. (2015). When applied to both pediatric and adult cohorts ALLSorts was able to classify previously undefined samples. ALLSorts is open source and publicly available to aid in further utilising RNA-seq data as a modality to investigate B-ALL.

## Methods

### Data sets

A combination of RNA-Seq, raw gene expression counts, and clinical information were obtained for 322 pediatric, 68 adult and 1988 mixed age B-ALL patients (Table 1). Furthermore, a dilution study undertaken by the Royal Children’s Hospital (RCH) was utilised to determine the effect of tumour purity (Table 1). In total, 2370 samples were available for use within this study. After gene expression counts were obtained for each dataset, gene identifiers were converted to symbols for consistency. Non-coding genes were discarded resulting in a final 20656 genes being included. The available samples were then split into a training set and series of test sets - summarised in Table 1 and detailed in Supplementary Table 1.

**Table 1.** Datasets used for training and validating the ALLSorts method. Datasets that are split across training and test are stratified by subtype. Note: Samples previously subtyped as “Other” or multi-label were removed in each set apart from the Multi-labels samples, which were tested separately. A breakdown of the samples used in the final training set are listed in Supplementary Table 1.

| Cohort | No. Samples | Train (%) | Test (%) | Purpose | Source |
| --- | --- | --- | --- | --- | --- |
| **St. Jude Children Hospital** | 1847 | 70 | 30 | Train & Hold out | (Gu et al., 2019) |
| **Lund University** | 195 | 70 | 30 | Train & Hold out | (Lilljebjörn et al., 2016) |
| **Royal Children's Hospital** | 127 | 0 | 100 | Paediatric | Partly Published (Brown et al., 2020) |
| **Peter MacCallum Cancer Centre** | 68 | 0 | 100 | Adult | Molecular Haematology Diagnostic Laboratory, Peter MacCallum Cancer Centre |
| **Royal Children's Hospital (dilution)** | 16 | 0 | 100 | Purity | (Brown et al., 2020) |
| **St. Jude Children Hospital and Lund (Multi-label)** | 117 | 0 | 100 | Multiple Subtypes | (Gu et al., 2019) |

### Data processing and feature creation

Training was broken down into four key steps: Preprocessing, Feature Creation, Standardisation, and Model Creation. These were encapsulated within a 10-fold cross validation (with replacement) and a nested grid search for optimal hyperparameter selection (Cawley & Talbot, 2010). Preprocessing first involves filtering for genes with a minimum of 10 counts in as many samples as the subtype with the lowest membership. This has previously been suggested as an initial step for differential gene expression analysis to remove low information genes (Chen, Lun, & Smyth, 2016). The filtered raw counts are then transformed to log2 counts per million (CPM) to scale samples by library size. Further scaling is then applied using the factors calculated from the Trimmed Mean of M-values (TMM) method (Robinson & Oshlack, 2010).

The Feature Creation step then generates an additional five sets of features from the gene expression counts, representing: fusions, iAMP21’s expression motif, chromosome ploidy, the B-ALL immunophenotype, and distance to a subtypes centroid. The fusion features are designed to capture the relative change in gene expression between the fusion gene partners that define known subtypes (*BCR-ABL1*, *TCF3-PBX1*, *ETV6-RUNX1*, *TCF3-HLF*), calculated by taking the difference in log_2_(CPM) between the 5’ and 3’ genes. Secondly, iAMP21 specific features are also created to reflect its distinct expression pattern across chromosome 21 (Tsuchiya et al., 2017). One region (region 3) is observed to be the most amplified so the feature takes the median expression of this region and compares it to the flanking regions using a log ratio. This is visualised in Supplementary Figure 1. This feature is added to ALLSorts along with the median value of each of the iAMP21 regions. Thirdly, a set of features that represent each chromosome’s relative expression is created. First, for each gene in the cohort, the median absolute deviation (MAD) is calculated, a median filter of five consecutive genes is applied, and finally taking the median of these values per chromosome. Fourth, a feature is also generated that attempts to quantify the immunophenotype of malignant B cells. As such, genes which are characteristic of B-ALL (*CD19*, *CD34*, *CD22*, *DNTT*, and *CD79A*) were chosen from the literature and have their preprocessed counts summed per sample to represent a B-ALL feature (Cobaleda & Sánchez-García, 2009). Fifth and finally, in an attempt to capture non-linear relationships associated with a subtype, a feature that represents the euclidean distance towards a subtype’s centroid in a nonlinear projection is calculated for each sample (Supplementary Method).

A standardisation step is then applied to the preprocessed counts and the new features to create a consistent scale across features, aiding in a model’s interpretability when assessing coefficients. This is achieved by calculating the z-score feature-wise.

### Training and running ALLsorts

The resulting counts matrix was input into a hierarchically organised set of logistic regression classifiers which were trained using the One Versus Rest method (Pedregosa et al., 2011). The logistic regression classifier includes an integrated feature selection method, L1 Regularisation, which shrinks coefficients of weak features to 0 - effectively negating their influence (Pedregosa et al., 2011). This can be seen as an embedded, multivariate feature selection step. During cross-validation, the most optimal hyperparameters and subtype probability thresholds are chosen. Hyperparameters are options that pertain to the learning algorithm itself and are not learned through exposure to data, e.g. the strength of the regularisation. Subtype probability thresholds are chosen by selecting a cutoff that maximises the F1 score of a fold and then averaging the result over all folds. Further details and descriptions of all the steps and the variables used can be found in the Supplementary Method.

The ALLSorts classifier was trained on a B-ALL cohort consisting of 1223 samples from two sources. While the pre-trained classifier is provided as part of our software package, users may also run the training stage themselves for their own projects if desired.

Applying the pre-trained model to new samples follows a similar sequence as described above. The key difference, however, is that each sample is normalised, standardised, and has custom features constructed according to the parameters chosen during training. The sample is then input into the hierarchy of logistic regression classifiers and probabilities are calculated individually, one versus rest, for each of the subtypes. Probabilities for the subtypes nested within a meta-subtype have their probabilities multiplied by the probability of that meta-subtype. Where any subtype’s probability exceeds the threshold, ALLSorts classifies the sample accordingly. This process is depicted in Figure 1.

**Figure 1.**
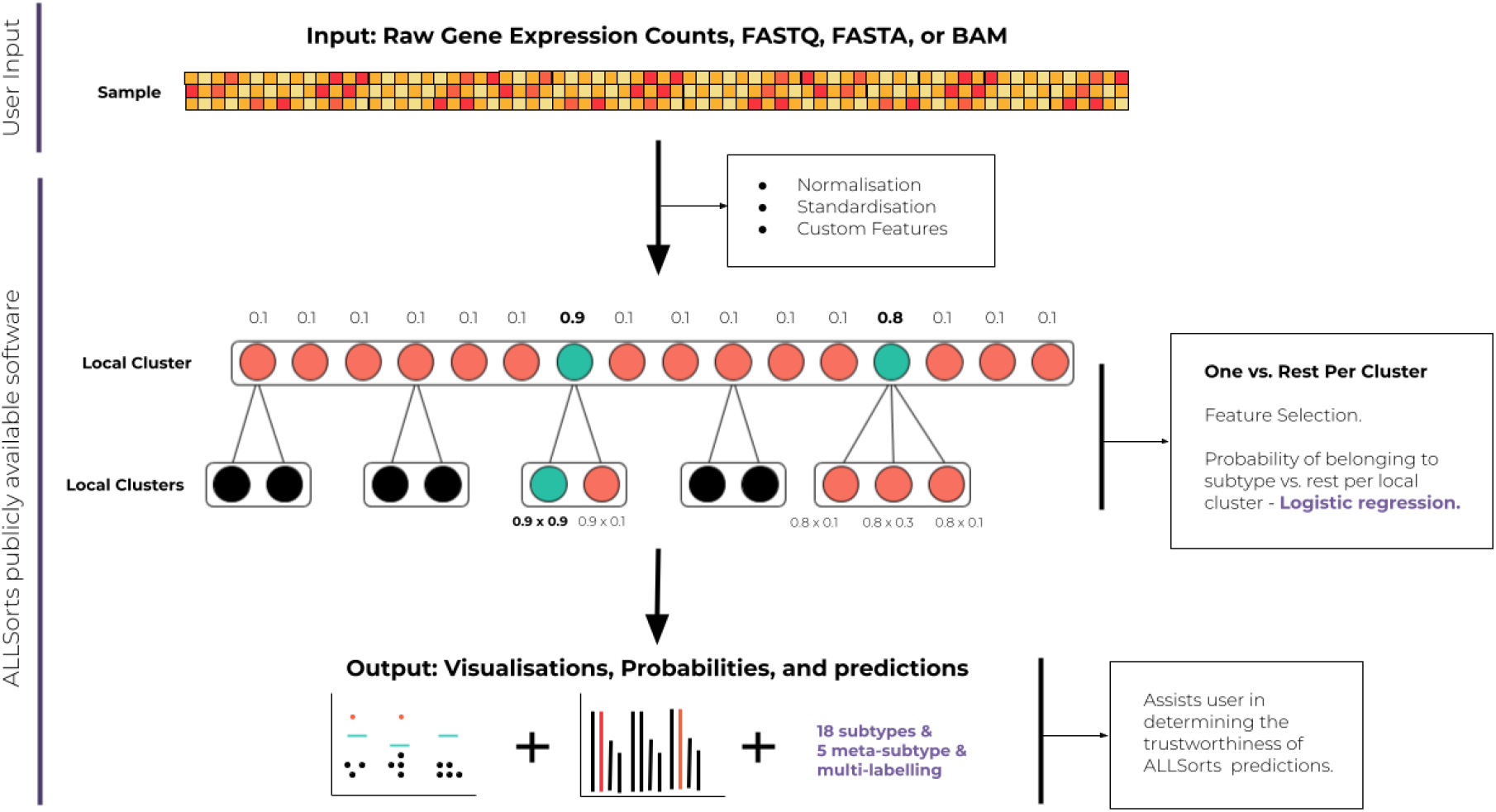
Overview of the ALLSorts classification strategy for new input. Green circles are where the probability exceeds threshold. No probabilities are calculated for the black circles as classification terminates at their meta-subtypes. In this example, two meta-subtypes exceed their thresholds at the first level. However, only one nested subtype succeeds. This would result in a multi-label classification consisting of the deepest subtypes/meta-subtypes that exceeded their respective thresholds.

## Results

### The ALLSorts Classifier

ALLSorts is a publicly available, pre-trained RNA-Seq classifier that attributes B-ALL samples to 18 known subtypes as depicted in Figure 2. This tool accepts one of three starting inputs: FASTQ/FASTA, BAM, or a matrix of raw counts for each gene in each sample to be classified. Input FASTQ/FASTA are aligned with STAR (Dobin et al., 2013). Resulting or supplied BAM files are gene level summarisation by featureCounts (Liao, Smyth, & Shi, 2014). This results in a consistent gene expression matrix for input into ALLSorts. The gene expression matrix then undergoes preprocessing, feature creation, and is filtered for the selected features as defined in Methods. The transformed counts are then input to the hierarchically organised logistic regression classifiers. The outputs are a list of predicted subtypes for the sample and a table of probabilities of subtype and meta-subtype membership per sample. There are also two visualisations that help verify the validity of the prediction and explore unclassifiable samples. The first visualisation depicts each user submitted sample’s probability of being a subtype relative to the determined threshold (e.g. Figure 3A). The second visualisation, termed waterfall plots, compares the maximum subtype probability for each sample to the probabilities of samples known to belong to that subtype (e.g. Figure 3B). The comparison samples shown by ALLSorts are from the held-out cohort, with 10 random samples selected per subtype (where available). These are packaged within the ALLSorts software but users can generate their own as described in the softwares documentation.

**Figure 2.**
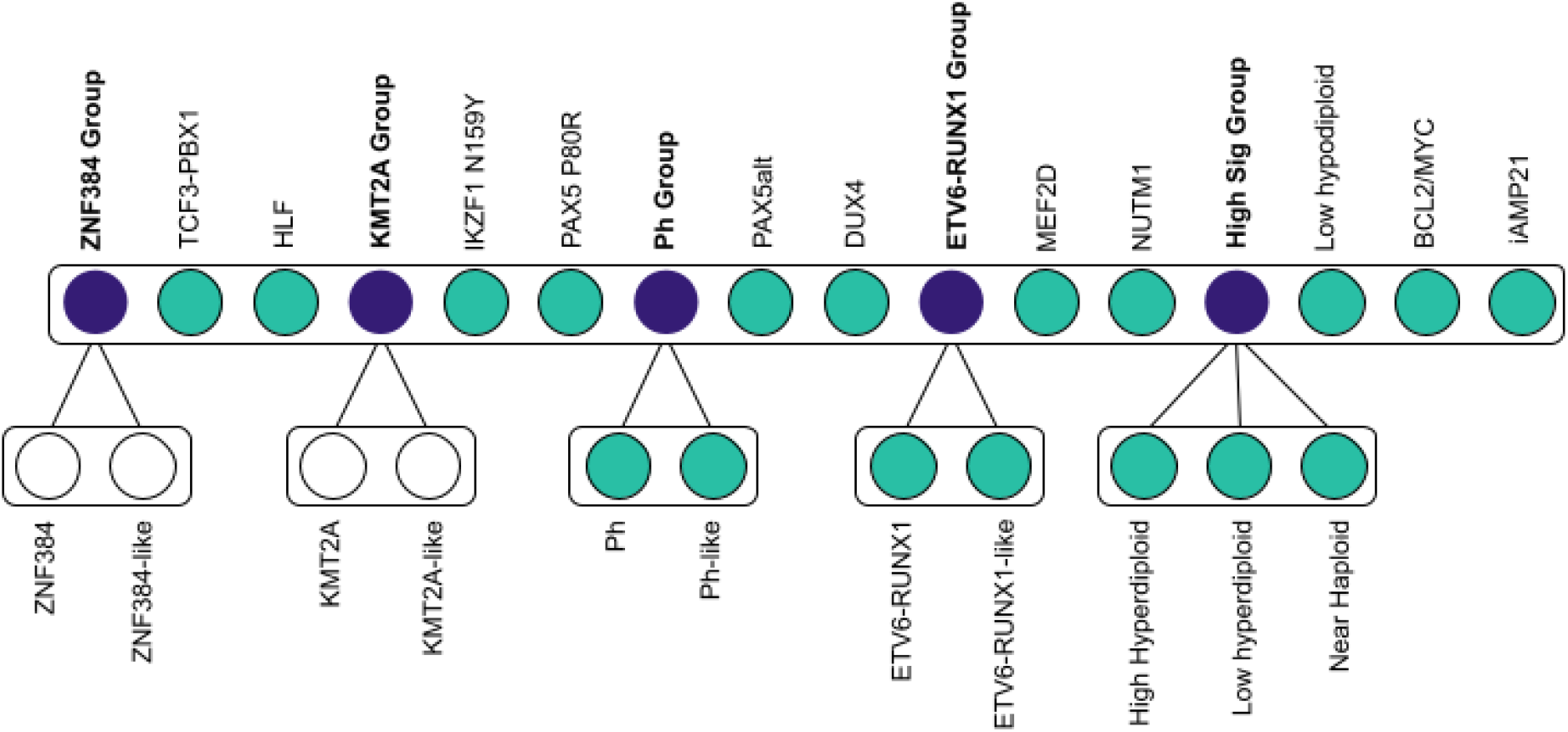
Overview of the ALLSorts classification architecture. In purple are the meta-subtypes that represent classes that have convergent or overlapping signals as per its nested subtypes. Green nodes are terminal subtypes. White nodes exist in the hierarchy, but classification currently terminates at the parent node due to a lack of training samples. CRLF2(Non Ph-like) is not included in this classification as its identification is better suited to downstream analysis. IGH-IL3 is also not included given only a single case across all cohorts.

**Figure 3.**
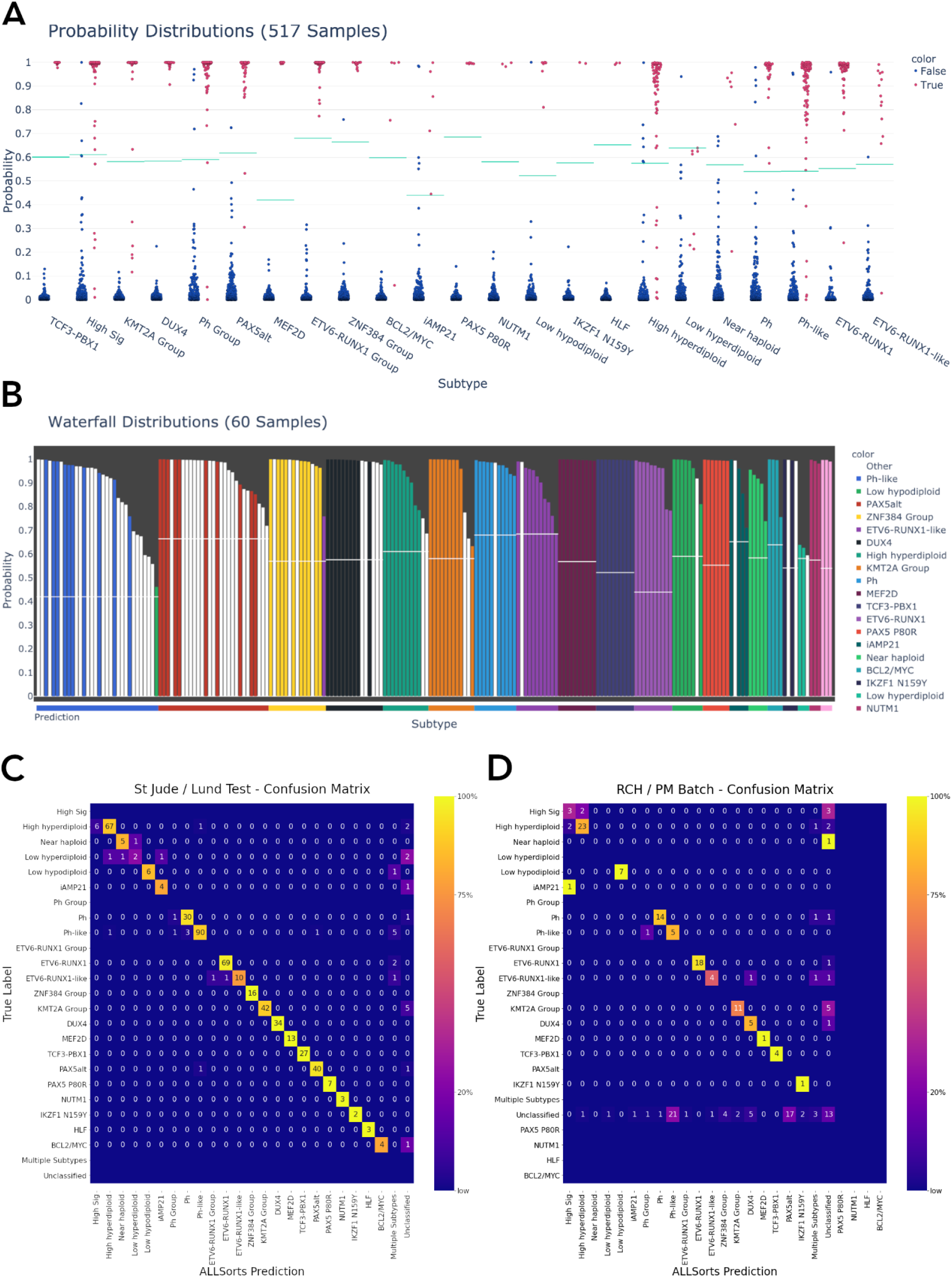
**A.** Probability distributions of St. Jude’s and Lund hold-out samples per subtype. For each sample, AllSorts reports a probability for every subtype. Blue dots are samples negative for that subtype, red are positive. The green lines are the subtype probability thresholds which were determined from the training data (Methods). **B.** Waterfall plot of RCH and PM samples that were previously Unclassified and then assigned subtypes by ALLSorts (white bars). The coloured bars represent samples with positive classifications from the St. Jude’s/Lund held out set. The Y axis shows the highest probability reported for any subtype, the X axis colour is the prediction as made by ALLSorts. Samples with multiple subtypes are displayed in every subtype where a prediction is made. **C.** Confusion Matrix of the St. Jude’s / Lund held-out test data. A confusion matrix shows the performance of the classifier. The Y axis represents the ground truth of each subtype, the X axis is the ALLSorts prediction. A perfect classification result would include no values off the diagonal. ALLSorts can predict samples to have multiple labels, these are reflected in the ‘Multiple Subtypes’ category. ‘Unclassified’ is where a sample’s probability did not exceed the threshold for any subtype. **D.** Confusion Matrix of the combined RCH and PM cohorts. The Y axis represents the previous classification of each sample, the X axis is the ALLSorts prediction. Rows without values indicate no subtype with that true label in the dataset.

### ALLSorts has biology inspired architecture using meta-subtypes

Phenocopies, where subtypes share a common transcriptional profile despite having different causal lesions, are a phenomenon in B-ALL (Li et al., 2018; Mullighan, 2019). ALLSorts represents phenocopies through the multi-level architecture depicted in Figure 2, where phenocopies share a meta-subtype that captures their shared transcriptional profiles. The classifier will first determine a sample’s meta-subtype and then undertake a more focussed classification between the nested subtypes.

There are five meta-subtypes that ALLSorts considers: ZNF384 Group, KMT2A Group, Ph Group, ETV6-RUNX1 Group, and High Ploidy Signature Group (High Sig). ZNF384, KMT2A, Ph, and ETV6-RUNX1 groups are nested with their phenocopy counterparts, (*-like)* subtypes (Figure 2). In addition, a High Sig meta-subtype consists of all subtypes that share a similar gene expression profile to High hyperdiploid. This includes Near haploid, Low hyperdiploid, and High hyperdiploid. Though Near haploid membership into this group would seem counter-intuitive, this subtype is considered to commonly harbour both diploid and hyperdiploid clones (Harrison et al., 2004; Safavi et al., 2013). In addition, as gene expression is a relative measure between genes, the expression of a haploid or diploid sample is potentially difficult to distinguish using gene expression counts.

### ALLSorts classifies 18 B-ALL subtypes

ALLSorts classifies 18 of the 23 subtypes recently described in Gu et al, (Gu et al., 2019). The CRLF2 subtype, seen by Gu et al., was removed as it lacked a distinct signal that could delineate it from Ph-like using gene expression data alone. The ZNF384-like and KMT2A-like subtypes seen by Gu et al. (2019) contained too few training samples to confidently train a discriminator. These two subtypes were included in meta-subtypes with their phenotypic counterparts and classification is terminated at the meta-subtype (Figure 2).

### ALLSorts is accurate on held-out validation data

To validate the generalisability of ALLSorts in classifying subtypes, 10-fold cross-validation was undertaken within the training set. In addition, the trained classifier was also applied to held-out test sets, which were split from the cohorts prior to training (Table 1). Four standard statistical metrics were used in the evaluation of the classifier: Accuracy, Precision, Recall, and F1 Score. Accuracy is the proportion of samples that were predicted correctly. Precision and recall are complementary, measuring the proportion of true positives and false negatives, respectively. Finally, the F1 score reflects the balance between precision and recall. These are calculated for each subtype and then aggregated by weighting the proportion of samples in each subtype.

Having both the cross-validation and held-out test set results from the same cohorts used for training allows us to determine whether the model is underfit or overfit. Table 2 demonstrates that the held-out test set has a higher precision, recall, F1 score, and a slightly lower accuracy than the cross-validation result. The confusion matrix of the held-out test set shows there is an imbalance of subtype classification performance (Figure 3C) with the highest levels of misclassification occurring between subtypes within the high (ploidy) signature group.

**Table 2.** ALLSorts performance in 10 fold cross validation determined during training and performance of held-out test sets from the St. Jude’s and Lund cohorts.

| Dataset | Accuracy (%) | Precision (%) | Recall (%) | F1 (%) |
| --- | --- | --- | --- | --- |
|  |  | Weighted average |  |  |
| Cross-Validation<br>(avg over 10 fold) | 90 | 96 | 91 | 93 |
| St. Jude's & Lund<br>hold out | 92 | 97 | 92 | 94 |

The predicted probability distributions for each sample in each subtype are plotted in Figure 3A which shows a dichotomy between subtypes that are clearly defined, such as DUX4 and TCF3-PBX1, compared with those with more of a continuum, such as High hyperdiploid. The subtypes that show well defined groupings are classified with better accuracy and are often heavily weighted by a single feature, typically expression of one of the fusion partners e.g. *PBX1* in *TCF3-PBX1*, *HLF* in *TCF3-HLF*, and *NUTM1* in its respective subtype. Those subtypes with a larger spread of probabilities are defined by larger collections of genes/features, such as aneuploidies, which tend to be driven by a wide range of genomic aberrations which and, produce more varied patterns of gene expression that impedes classification (Gu et al., 2019).

Within the held-out dataset results, only two of seven Low hyperdiploid samples were correctly called (Figure 3C). Of the five failed calls, two were called as Unclassified and three misclassified as iAMP21, Near haploid, and High hyperdiploid. Near haploid saw a misclassification as Low hyperdiploid. This is not unexpected, given it is common for Near haploid samples to have both Diploid and High hyperdiploid clones. Finally, High hyperdiploid had six misclassifications deferring to the parent meta-subtype, one to Ph-like, and two being marked as Unclassified. Therefore, given the ambiguity of the High Ploidy Signature subtypes, one can conclude that further data is required to extract a common signal or a new approach is required. In the interim, the High Ploidy Signature meta-subtype can be used with an accuracy of 93%. In addition, both Ph/Ph-like and ETV6-RUNX1/ETV6-RUNX1-like saw misclassifications to their phenotypic counterparts or to the meta-subtype (Figure 3C). In circumstances such as this, the ambiguity can be resolved by running a fusion finder on these samples post classification and filtering the output for fusions known to drive these subtypes.

### ALLSorts applied to independent cohorts

Although the results on the held out data show that the ALLSorts classifier performs well across most subtypes, the test dataset was from the same cohort as the training dataset. Typically, ALLSorts will be applied to new samples which include technical differences in the acquisition and processing of the samples compared to the training data. To test whether ALLSorts is robust to such effects we applied it to paediatric and adult B-ALL cohorts from the Royal Children’s Hospital (RCH) and Peter MacCallum Cancer Centre (PM), respectively. Each cohort had different sequencing and library preparation protocols making them an effective representation of a typical input.

ALLSorts predictions of the combined cohort are depicted within the confusion matrix in Figure 3D. To evaluate these results they can be broken into four categories: match with ground truth, new classification into a subtype, reclassification to another subtype(s), and subtype to unclassified. Assuming the matched samples are correct (109 samples or 56%), only samples described by the latter three categories required further exploration.

Of the 74 samples that were previously Unclassified, 61 (82%) were newly classified into one of the 18 subtypes or five meta-subtypes offered by ALLSorts. Of these, 42 were evaluated to be plausible, one was incorrect (tumour purity 12%), and no definitive evidence could be made for 17. Curiously, five of the samples without suitable evidence were classified as DUX4 despite the fusion callers, JAFFA and Arriba, not finding a relevant fusion gene (Supplementary Table 1).

Reclassification to a new subtype accounted for 10 samples. Of these, eight were correct if including the meta-subtype as a positive call. One sample was incorrectly called High Sig instead of iAMP21. However, this sample had a tumour purity of only 13% which could account for this misclassification. Finally, one contained a novel *ETV6* fusion but was predicted as being DUX4. The reason for this is currently unknown.

The most important misclassifications to explore were the 15 samples (7.7%) previously labelled as a distinct subtype which ALLSorts assigned as Unclassified. Six of these samples had tumour purities of less than 10%, which may account for misclassifications in these cases, i.e. ALLSorts was found to be adversely affected by tumour purities under this (Supp Figure X). Of the remaining nine, three were previously labelled as KMT2A rearranged of which each had cytogenetic evidence of the relevant fusion genes. However, each of these samples exhibited low expression for genes such as *MEIS1,* a typical target of KMT2A fusions which ALLSorts weights highly in KMT2A Group classification. Four High Sig samples with a tumour purity above 10% did not reclassify according to their ground truth. However, two had high probabilities of being ETV6-RUNX1 Group and each had an associated *ETV6-BCL2L14* fusion discovered through Arriba. As these two samples also had relatively high probabilities for High Sig (over 39%), it is possible that this is a multi-subtype sample. The remaining newly Unclassified samples were labeled according to cytogenetics only or had a lower tumour purity (~16%).

A full list of samples that had unexpected classifications with potential causative variants found is available in Supplementary Table 3. Of these 86 samples, 56% had a plausible explanation that the ALLSorts classification was correct at least to the meta-subtype level, 8% were incorrect, 27% remained ambiguous in terms of evidence supporting or dismissing plausibility of the call, and 9% were defined as having tumour purity too low for concrete classification (less than 10%). Summary statistics with adjusted labels can be seen in Table 3.

**Table 3.** ALLSorts performance in the RCH and PM cohorts once orthogonal evidence gave plausibility to the calls. Two sets of summary statistics are presented, representing where ambiguous samples have been marked as correct or as incorrect - indicating the boundaries of the classifiers performance on these datasets.

| Ambiguous Samples | Accuracy (%) | Precision (%) | Recall (%) | F1 (%) |
| --- | --- | --- | --- | --- |
|  |  | Weighted average |  |  |
| Marked as incorrect | 82 | 92 | 94 | 92 |
| Marked as correct | 93 | 99 | 97 | 98 |

### ALLSorts classifies samples with multiple subtypes

One unique feature of ALLSorts is its ability to classify samples into more than one subtype. The St. Jude and Lund cohorts included 117 samples that were described as having multiple subtypes and were not excluded based on the sample filtering outlined in Methods (Table 1). Both subtypes were based on both gene expression analysis and cytogenetics. Without specifically training ALLSorts to recognise samples exhibiting multiple subtypes, this cohort was used to investigate the capacity for multi-label classification.

We found the probability of getting at least a single subtype correct is 86.31% and 90.5% if including meta-subtypes (Table 4). Given this is similar accuracy to the single subtype benchmarks, multi-label classification can be added without reducing single subtype classification accuracy (Table 2). However, we only predicted both subtypes 26% of the time. Interestingly, within the held-out test set thought to be composed of samples with only a single subtype, nine samples were predicted as having two. Similarly, the PM and RCH combined cohort had six samples (3%) classified with two subtypes. Of these six samples five were found to have evidence pointing to the accuracy of both calls from fusion calling and karyotyping (Supplementary Table 3). This demonstrates that multiple label classification with ALLSorts can add further value of a classifier with little cost in performance. In future, as further manual labelling of multi-label samples becomes available for use in training data, these multi-label subtypes could be explicitly trained for.

**Table 4.**
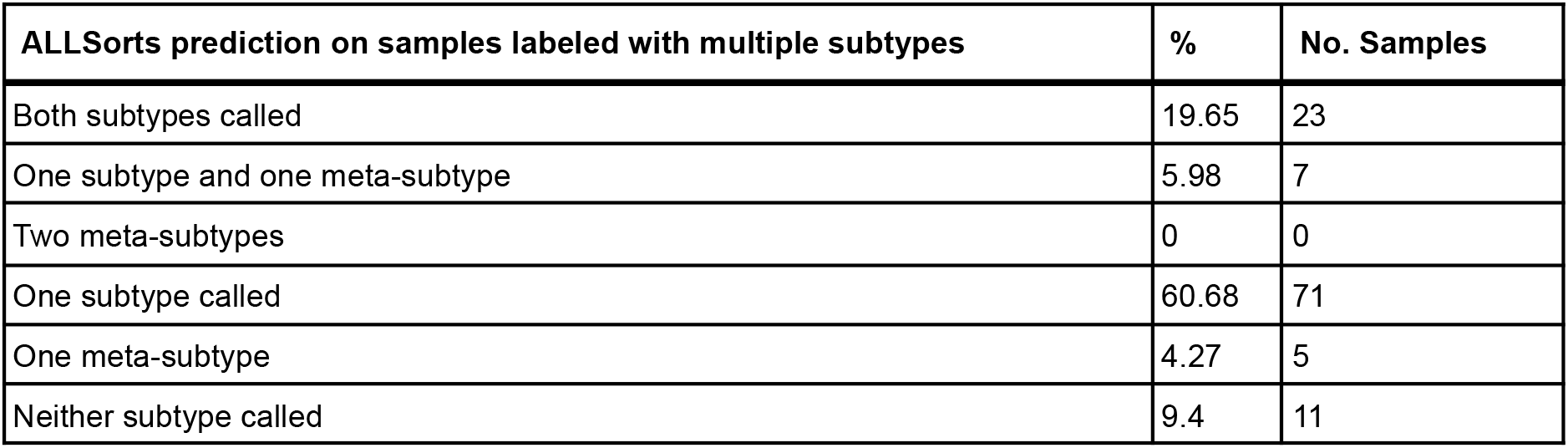
Breakdown of multi-label predictions.

| **ALLSorts prediction on samples labeled with multiple subtypes** | **%** | **No. Samples** |
| --- | --- | --- |
| Both subtypes called | 19.65 | 23 |
| One subtype and one meta-subtype | 5.98 | 7 |
| Two meta-subtypes | 0 | 0 |
| One subtype called | 60.68 | 71 |
| One meta-subtype | 4.27 | 5 |
| Neither subtype called | 9.4 | 11 |

## Discussion

In this study we present ALLSorts, a B-ALL subtype classification tool that can precisely attribute samples to 18 subtypes and five meta-subtypes according to their gene expression measurements. This tool has been trained and validated with a combined cohort of over 2300 samples and is offered for public use through Github. Minimal setup is required to prepare the input and users receive subtype probabilities, predictions, and visualisations as output.

One of the novel contributions of ALLSorts is a hierarchical architecture representative of subtypes, their phenocopies, and the broader encompassing signal defined as meta-subtypes. There are clear clinical advantages of such an approach. In an ideal scenario, a classification algorithm can attribute a sample to a subtype with 100% accuracy. However, in the case where a sample cannot be confidently ascribed it may still be clinically useful to classify in a higher tier (Galea et al. 2017). For example, if it is difficult to delineate between Ph and Ph-like, a positive Ph Group classification can be further investigated for specific targetable lesions. There are also technical advantages of hierarchical gene expression classification. With a growing number of subtypes the ability for a classifier to discriminate becomes more challenging. Segmenting the classification process into a hierarchy can alleviate this and result in higher accuracy (Galea et al. 2017; Silla and Freitas 2011). Indeed, in our testing, the hierarchical architecture (Figure 2) demonstrated an improvement in performance over a flat architecture during the 10 fold cross-validation of ALLSorts (accuracy of hierarchical architecture = 87.1%, accuracy of flat architecture = 83.6% - Supplementary Table 2). With greater understanding of the underlying mechanisms of tumorigenesis, more nuanced hierarchies may be proposed and further improve both interpretability and performance of subtype classification models.

A key component to this study was testing the software across cohorts with various biases to verify its robustness. Comparing the hold-out and validation confusion matrices in Figure 3, on first look, suggests ALLSorts performance on the validation set has some inaccuracies. There are multiple reasons as to why this may be the case. First, the subtype schema used to identify the previous subtypes in the combined cohort was limited to nine in total: Ph, Ph-like, Hyperdiploid, Hypodiploid, ETV6-RUNX1, ETV6-RUNX1-like, TCF3-PBX1, DUX4, KMT2A, and Unclassified. Therefore, we expect the subtype designation to change in some cases as samples begin to classify according to the new schema. Secondly, previous classifications were derived through a combination of bioinformatics tools (fusion finding and classification) and cytogenetic methods (G-Banding Karyotyping and Fluorescent in situ hybridization). Brown et al (2020) demonstrated that RNA-Seq, despite its advantages, does have limitations that can be overcome through precise cytogenetic methods. This is perhaps shown in the *KMT2A* samples that were attributed Unclassified by ALLSorts, as *KMT2A* rearrangements are known to be lowly expressed and evade multiple bioinformatics methods (Brown et al., 2020). However, the converse is also true and ALLSorts was capable of classifying samples where the driving event was not readily clear through traditional cytogenetics. For example, two plausible ETV6-RUNX1 Group classifications were made from samples that were previously not captured through G-Banding or FISH. Regardless, in many of the cases with apparent misclassifications there was some justification for the ALLSorts call based on orthogonal information from the sample or patient (Supplementary Table 3). Given the summary statistics outlined in Table 3 accounting for these justifications, we believe ALLSorts is accurate and has the ability to provide new annotation even in previous B-ALL cohorts.

As the number of B-ALL samples that are sequenced increases there will be improvements that can be made to the classifier. ALLSorts has the ability to retrain the classifier as more samples become available. The most obvious improvements will be retraining for classification of subtypes that have relatively low numbers of samples, such as BCL2/MYC. In addition, although gene counts are clearly useful in determining the overall patterns of expression in a subtype, a more refined method that uses more nuanced aspects of the data such as transcript quantification, equivalence classes, or kmers could provide increased performance.

In summary, ALLSorts is an accurate, comprehensive and freely available classification tool for determining subtypes of B-ALL.

## Supporting information

Supplementary Results

## Acknowledgements

- Tumour samples and coded data were supplied by the Children’s Cancer Centre Tissue Bank at the Murdoch Children’s Research Institute and The Royal Children’s Hospital (https://www.mcri.edu.au/childrenscancercentretissuebank). Establishment and running of the Children’s Cancer Centre Tissue Bank is made possible through generous support by CIKA (Cancer In Kids @ RCH), The Royal Children’s Hospital Foundation and the Murdoch Children’s Research Institute.
- Raw gene expression counts for B-ALL tumour samples used for analysis in this study were obtained from St. Jude Cloud (https://www.stjude.cloud) – a publicly accessible pediatric genomic data resource requiring approval for controlled data access.
- This work was supported by grants from the Wilson Centre for Lymphoma Genomics and the Snowdome Foundation.
- This work was funded by NHMRC project grant APP1140626.

## Authorship Contributions

BS: Conceptualization, Formal Analysis, Methodology, Software, Visualisation, Writing – Original Draft Preparation, Writing – Review & Editing; AO: Conceptualization, Supervision, Methodology, Writing – Original Draft Preparation, Writing – Review & Editing; NMD: Conceptualization, Supervision, Methodology, Writing – Original Draft Preparation, Writing – Review & Editing; LB: Clinical Expertise, Writing – Review & Editing; GR: Samples Supply, Clinical Expertise, Writing – Review & Editing. AL: Bioinformatics Support, Writing – Review & Editing. HK: Biological Expertise, Writing – Review & Editing. LL: Sourcing orthogonal clinical information, Writing – Review & Editing. IM: Biological Expertise, Writing – Review & Editing. PB: Clinical Expertise, Writing – Review & Editing. PE: Clinical Expertise, Writing – Review & Editing.

