## Supplementary Figures for "ALLSorts: a RNA-Seq classifier for B-Cell Acute Lymphoblastic Leukemia"

### Supplementary Figure 1. iAMP21 Motif

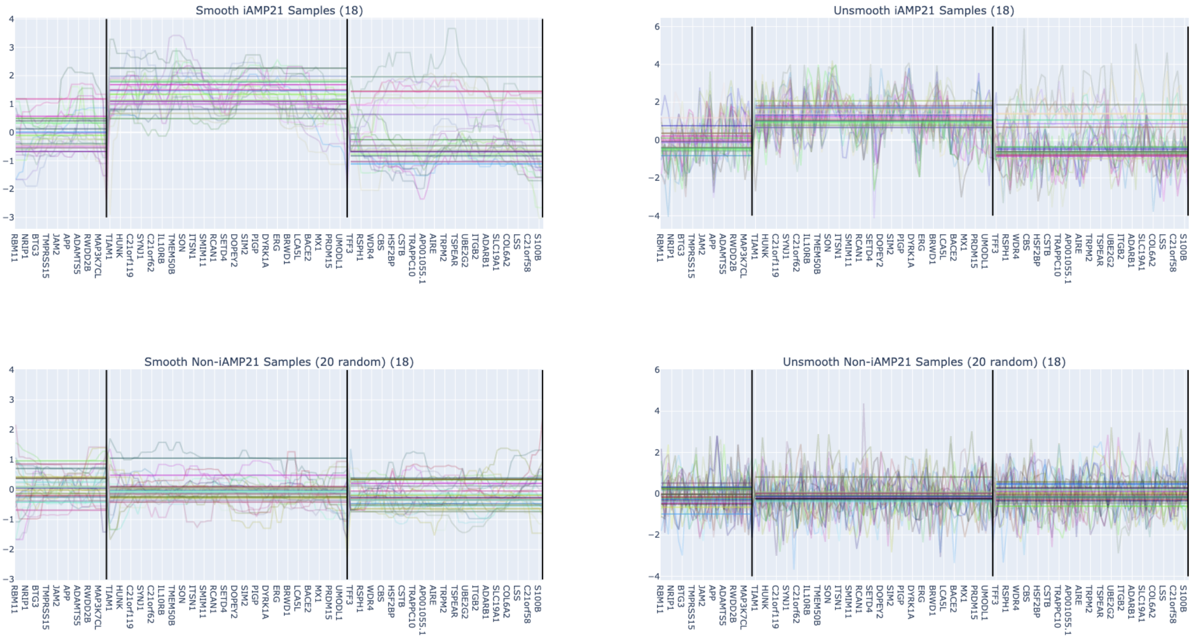


Supplementary Figure 1. iAMP21 as depicted by the ordered genes in chromosome 21 is divided over four bins (first bin contains no genes filtered through ALLSorts pre-processing). Samples with confirmed iAMP21 (top) vs normal (bottom). Left plots use iterative median smoothing, right does not. A clear motif is apparent between those samples with iAMP21 and those without.

### Supplementary Figure 2. Ploidy Features

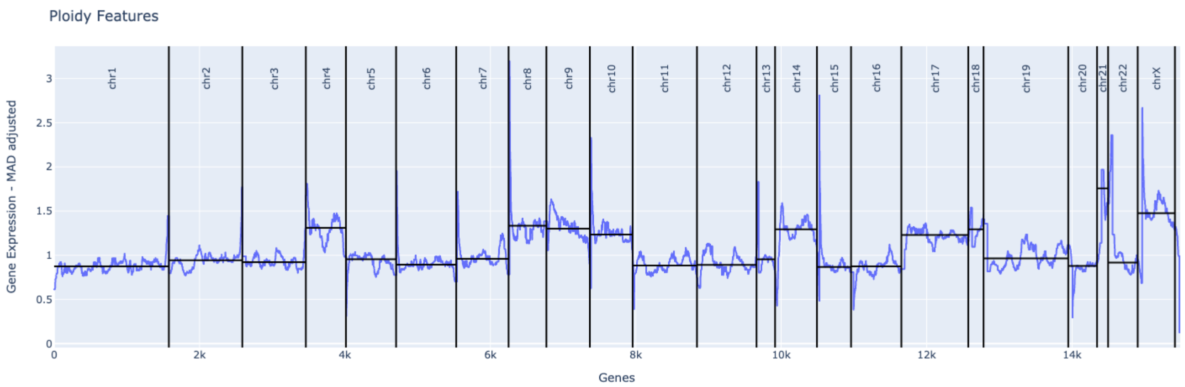

Supplementary Figure 2. St. Jude Children's Research Hospital sample with ***'56,XX,+X,+4,+6,+8,+10,+14,+17,+18,+21,+21[18]/46,XX[2]'*** karyotype. Genes are ordered according to their position on each chromosome, median absolute deviation is calculated across all samples in the training set per gene, then median iterative filtering is applied. Finally, a median across each chromosome is calculated and is used as a feature within the model.
