## Supplementary Results for "ALLSorts: a RNA-Seq classifier for B-Cell Acute Lymphoblastic Leukemia"

### Supplementary Results 1. ALLSorts is robust to low tumour purity

To test the effect of tumour purity on classification performance, two dilution series were used. These were created from a mixture of RNA extracted from the lymphoblastoid cell line from NA12878 with either a Ph (*BCR-ABL1* fusion) tumour sample or a Ph-like (*PAX5-JAK2* fusion) tumour sample (Brown et al., 2020). The tumour RNA was mixed in the proportions: 0%, 10%, 20%, 30%, 40%, 60%, 80% and 100% and each mixture was classified with ALLSorts. As expected, purer tumour samples corresponded to higher probabilities (Figure 4). For the Ph+ dilution series ALLsorts was able to classify the correct subtype down to a purity of 10%. The Ph-like dilution series classified correctly down to 20% purity and at 10% deferred to being classified as Ph Group. Though a higher tumour purity will result in more confident classifications, these results suggest ALLSorts is robust to tumour proportions of above 20% in these subtypes. However, this would need to be tested for each subtypes in order for a claim of general robustness to tumour purity could be made.

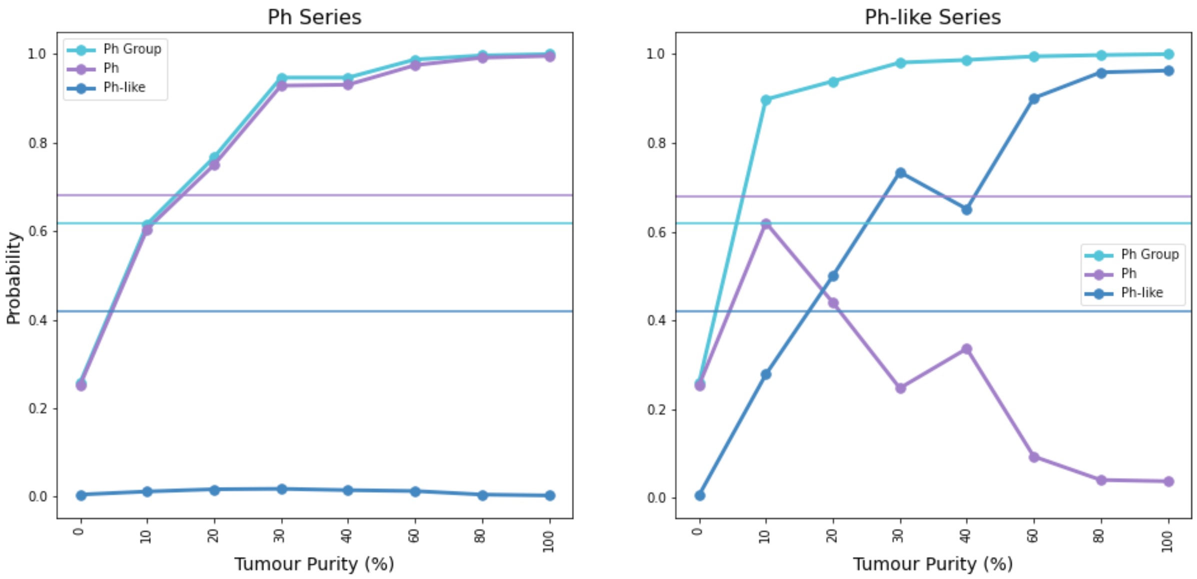


Figure 4. Probabilities from ALLsorts of the Ph Group meta subtype and Ph and Ph-like subtypes in response to tumour dilution. Subtype probability thresholds are indicated by the horizontal lines.

To visualise this within the combined cohort, samples with available blast percentage within both the PM and RCH cohorts were plotted against the plausible, ambiguous, and incorrect categories discussed in Results (Figure 5). Seven samples had a blast percentage of less than 30%. The single plausible sample was called a PAX5alt. Three ambiguous samples were classified as Ph-like, of which one was also called High hyperdiploid. The three incorrect classifications were: a DUX4 (IGH-DUX4 fusion) that was unclassified, a sample with normal cytogenetics called High hyperdiploid, and an iAMP21 misclassified as High hyperdiploid.

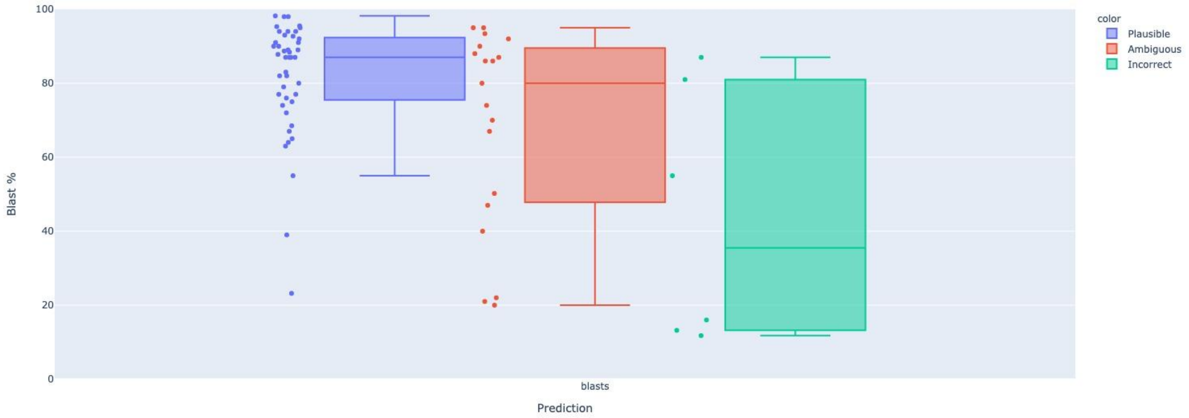


Figure 5. Blast % versus Plausible/Ambiguous/Incorrect categorisation allocated during investigation into known driver events.
