## Supplementary Method for "ALLSorts: a RNA-Seq classifier for B-Cell Acute Lymphoblastic Leukemia"

### Data used in this study

Gene expression counts with complimenting clinical information were obtained/created for four B-ALL cohorts and a dilution series. Raw counts for 1988 samples from a recent St. Jude Children's Research Hospital study were available for public download through the St. Jude Cloud's visualisation community (Gu et al., 2019; McLeod et al., 2021). Raw sequencing reads from 195 samples were obtained from Lilljebjörn et al. (2016) (Lund - accession id: EGAD00001002112), 127 paediatric samples from the Children's Cancer Centre Tissue Bank at The Royal Children's Hospital (RCH), Melbourne, Australia (Brown et al., 2020) and 68 adult samples from the Molecular Haematology Laboratory, Peter MacCallum Cancer Centre, Melbourne, Australia (PM). The Lund, RCH, and PM datasets were mapped to the human reference genome version hg19 using STAR 2.7.3a_2020-01-23 in 2 pass mode with quantMode set to Gene, with otherwise default options. Gene expression counts for the RCH, Lund, and PM dataset were provided as output from STAR, using the GRCh37.87 annotation obtained from the ensembl FTP (ftp://ftp.ensembl.org/pub/grch37/current/gtf/homo_sapiens/Homo_sapiens.GRCh37.87.chr.gtf.gz). After gene expression counts were obtained for each dataset, Ensembl gene identifiers were converted to gene symbols. Identifiers with multiple copies had their counts combined. After this, genes other than those with biotypes of protein coding or recognised as Immunoglobulin (Ig) variable chain or T-cell receptor (TcR) genes were discarded. Resulting in a final 20652 of the 57773 original genes being used in the training data.

The St. Jude samples were labelled according to the 23 subtypes outlined in Gu et al. (Gu et al., 2019). However, the Lund samples were assigned labels according to the following classes: *High hyperdiploidy, ETV6-RUNX1*, *Ph-like*, *MLL*, *TCF3-PBX1*, *DUX4-rearranged*, *BCR-ABL1*, *dic(9;20)*, *ETV6-RUNX1-like*, *B-other with fusion*, *B-other*, *without fusion*, *Hypodiploid*, *Near Tetraploid*, and *iAMP21*. As karyotype data was also available, the aneuploid samples were distributed across *High hyperdiploid* (58) and *Low hypodiploid* (1) accordingly. *MLL*, *DUX4-rearranged,* and *BCR-ABL1* were renamed *KMT2A*, *DUX4*, and *Ph*, respectively, to reflect the St. Jude naming conventions where fusion information was available. Samples labelled *dic(9;20)* and *Other* were removed and explored using the trained classifier. Samples that were not labelled as having multiple subtypes, but showed signs according to associated clinical information (i.e. *iAMP21* being mentioned in karyotype but not mentioned as a subtype) were discarded from the training data. In addition, St. Jude labelled aneuploid samples that did not have concordance with the karyotype were also discarded. In all, 168 samples were marked for exclusion from the training data across both the St. Jude and Lund cohorts. Finally, training and test sets were segmented according to Table 1 and the training samples listed in Supplementary Table 1.

### Training

10-Fold Cross Validation was performed during training to score the method and select optimal thresholds, whilst 3-fold cross validation was implemented in an inner loop to select the hyperparameters of the model. The scores averaged across the outer loop were then used as a benchmark for comparison to other datasets to consider overfitting.

Each of the following stages was performed within each fold to prevent the leakage of data between the resulting training and validation splits.

#### Pre-Processing

The first step of pre-processing is to filter lowly expressed genes as they contribute little information about the biology fundamental to the classification problem. To achieve this, the method outlined in Chen et al. (2016) was adopted. That is, the training set was transformed into counts per million (CPM) prior to filtering. This is to ensure that genes that are lowly expressed due to smaller library sizes are not naively filtered. Genes are retained if there are at least 10/L (where L is the smallest library size) in at least as many samples as the subtype with the lowest membership. Once lowly expressed genes have been identified they are stored for later removal in new samples input into the classifier. The training data is then normalised using a Python implementation of the Trimmed Mean of M-values (TMM) normalization method (Robinson & Oshlack, 2010). This method is preferable over methods such as fragments per kilobase million (FPKM) as it accounts for library composition and is therefore suitable for inter-sample comparisons (Conesa et al., 2016). The reference used to calculate TMM scaling factors is then stored for later use when ALLsorts is applied to a new dataset. The filtered raw counts are then transformed to log2 counts per million (CPM) to scale samples by library size. Further scaling is then applied using the factors calculated from the Trimmed Mean of M-values (TMM) method.

#### Feature Creation

ALLSorts uses four sets of manually crafted features to represent the biology of B-ALL represented in the literature. The first is a set of features that represent known fusion genes that are highly relevant to some subtypes: *ETV6-RUNX1*, *BCR-ABL1*, *TCF3-PBX1*, and *TCF3-HLF*. The resulting features are simply the log difference between the two partner genes. This feature is important to include as it encapsulates the relationship within a single feature. The second set of features represent the relative expression of each chromosome per sample. This was calculated in a similar fashion to existing solutions to calculating ploidy from RNA-seq (Gu et al., 2019; Serin Harmanci, Harmanci, & Zhou, 2020). The genes in the training set are first scaled by median absolute deviation.  Median iterative filtering is then applied across genes in each chromosome, smoothing the signal across the chromosome. A final median is then selected per chromosome and a set of 27 features are created from this (chr 1-22, X, Y, median across the chromosomes, and two representing how many are highly expressed and how many are low). A visualisation of these features can be seen in Supplementary Figure 2. Thirdly, an iAMP21_ratio feature is also created based on the knowledge of its distinct signal across chromosome 21 (Tsuchiya et al., 2017). For each sample chromosome 21 is divided into four bins and is scaled by median absolute deviation (Supplementary Figure 1). Fourthly, in an attempt to capture non-linear relationships associated with a subtype, a feature that represents the euclidean distance towards a subtype's centroid in a nonlinear projection is calculated for each sample. Concretely, for each local cluster of subtypes (Figure 1) a random forest machine learning classifier is trained in a one-versus-rest fashion. From this, the top 20 features as measured by feature importance are attributed to each of the subtypes. A Kernel PCA projection is then created for each subtype using these genes, hopefully delineating between the subtype and the rest. The centroid is then calculated across each of the true subtype samples in this projection and the euclidean distance to this point is then calculated per sample. The final feature is constructed from the difference between the bin at the highest and lowest points. Finally, a single B-ALL feature is created as the log sum of CD19, CD34, CD22, DNTT, and CD79A. These genes are known markers for B-ALL in the literature (Chiaretti, Zini, & Bassan, 2014; Cobaleda & Sánchez-García, 2009). The purpose of this feature is not to be necessarily used in the following classification stage, but rather as a filter to remove false positives when classifying healthy samples or other cancers.

### Feature Standardisation

Features are standardised through transformation into a z-score, by subtracting the mean and dividing by the standard deviation in a feature-wise manner. The end result of this is a set of feature distributions, each with a mean of 0 and a standard deviation of 1. Standardisation was shown to perform higher when utilised within a hierarchical architecture (Supplementary Table 2). In addition, standardisation by z-score allows the coefficients within a linear model to be equally considered in terms of importance, that is, higher coefficients are relative to the corresponding features importance.

### Hierarchical Classification

Given the large set of input genes (~20000) relative to the number of training samples (~2000), a subset of genes needed to be selected to reduce dimensionality and the risk of overfitting of a trained model. During training, three feature selection methods were competed to find the best performing, each would then be followed by a further selection using L1 Regularisation embedded within liblinear's implementation of the logistic regression model, the core classification algorithm used within ALLSorts. L1 regularisation is used to encourage sparsity within the model by penalising uninformative genes by trending their coefficients to zero (Pedregosa et al., 2011). This can be considered a multivariate feature selection method and is tuned with the parameter C - lower values will result in more aggressive regularisation. The first competing method is a univariate statistical test, mutual information, that is calculated per subtype (Pedregosa et al., 2011). This list is then rank-ordered and subjected to a two standard deviation from the mean cutoff with the intention of completely omitting a set of uninformative genes, whilst retaining genes with high value. The other method used a L1 regularised logistic regression classifier to pre-train the model and select the genes useful for classification, passing them on to the final classifier. The final method has no feature selection prior to the classifier, depending only on the single regularisation method embedded in liblinear. The best combination of hyperparameter values was determined by minimizing log loss over 3 fold cross validation (Supplementary Table 2). Although more folds would have been ideal, there were too few samples in many of the subtypes. Balanced accuracy was used as the metric of success, as opposed to purely accuracy, as this accounts for the imbalance of samples allocated across subtypes. Balanced accuracy is calculated according to:

$$(1)\ balanced\ accuracy\ =\ \frac{1}{2}\left(\frac{TP}{FP+TP}+\ \frac{TP}{FP+TP}\right)$$

The winning feature selection method was the single L1 regularised logistic regression model, without prior feature selection. The results of hyperparameter selection can be seen in Supplementary Table 2.

An important aspect of classification is assigning probabilities to a discrete class. This is typically performed by setting a 50% threshold per subtype. Though this is appropriate for binary classification problems it is not necessarily appropriate for multi-class classification problems. In these cases, the probability is distributed amongst multiple classes and may not exceed 50% in any. Many algorithms therefore surpass this problem by choosing the maximum probability from the result. However, this is not appropriate for cases where there is a chance that a sample belongs to a new class entirely. To attempt to resolve this problem, ALLSorts determines the probability thresholds using the cross-validation results from the training data. Thresholds are determined for each subtype, using one of two methods. In the first case, if the positive and negative samples for that subtype separate cleanly on probability, i.e. the highest probability for a negative sample is lower than the lowest probability of a positive sample, the midpoint between these two points is chosen. However, if this is not the case and the positive and negative samples overlap in probability, a threshold that maximises the F1 score is chosen. The threshold for any child subtype is weighted by the parent threshold through multiplication.

### Prediction

To predict B-ALL subtype, raw gene expression counts are input into the trained ALLSorts algorithm, this data is pre-processed and has features created using values acquired through the training set. These processed counts are then filtered by the genes selected during the training of the algorithm. Finally, they are input into the hierarchical classifier and their probabilities of belonging to each subtype is calculated. Furthermore, children probabilities are the multiplication between the child probability and its parent. After this, ALLSorts has various visualisation and prediction methods that help users in understanding their results.
